# Estrogen-Deficiency Degrades Left Ventricular Diastolic Function and Energy Metabolism in Hypertensive Female Mice

**DOI:** 10.64898/2026.06.05.730534

**Authors:** Aijun Zhang, Shumin Li, Indira Vedula, Somik Chatterjee, Yang Wu, Jianhua Gu, George G. Rodney, Karla M. Kurrelmeyer, Heinrich Taegtmeyer, Dale J. Hamilton, Henry J. Pownall, Anisha A. Gupte

**Author notes:** **Corresponding author:** Henry J. Pownall, PhD, Charles W. Duncan Jr., Department of Medicine, Houston Methodist Hospital and HM Academic Institute, 6670 Bertner Ave, Mailstop R10-121, Houston TX 77030-2602.

## Abstract

**Aims:** Heart failure (HF) due to diastolic dysfunction (DD) but normal left ventricular (LV) ejection fraction (EF) is termed HF with preserved EF (HFpEF). Given its high prevalence in post-menopausal women, we hypothesized that 17β-estradiol (E2) function is mechanistically linked to DD HF and investigated E2-deficiency in the etiology of DD in a mouse model.

**Methods:** Female C57BL/6J mice were divided into one group with hypertension induced by N(ω)-nitro-L-arginine methyl ester (L) and cardiac pressure overload induced by angiotensin II (A), collectively inducing DD, and one group with sham treatment. Groups were subdivided to receive an ovariectomy (OVX) to induce estrogen-deficiency and emulate menopause or sham surgery. During the next 21 days, mice were tested for cardiac function, food intake, response to E2 agonists, gene profiling, cardiomyocyte-contractility and elasticity, and cardiac mitochondrial function.

**Results:** OVX-associated E2-deficiency exacerbated DD in a time-dependent way without affecting EF or stroke volume, emulating a severe DD phenotype. These changes paralleled those for mitochondrial dysfunction, i.e., upregulation of genes associated with stress, energy metabolism, and fibrosis, as well as functional and structural defects in cardiomyocytes. Treatment of OVX + (L + A) with E2 or a G-protein-coupled estrogen receptor agonist normalized diastolic function, whereas estrogen receptor beta agonists did not. The OVX DD mice exhibited moderately impaired mitochondrial function, which delayed cardiomyocyte relaxation but not contraction, altered cardiac substrate utilization, reduced cardiomyocyte elasticity, increased production of reactive oxygen species, and potentiated extracellular fibrosis.

**Conclusions:** OVX-induced E2-deficiency generates metabolic, structural, and functional changes in cardiomyocytes and the adjacent extracellular matrix, exacerbating the effects of L and A on diastolic function. This robust DD model revealed a role for E2 via ERα in diastole-regulation in female mice and raised questions about similar mechanisms operative in postmenopausal women.

**HIGHLIGHTS:**

1. Ovariectomy (OVX)-induced estrogen-deficiency exacerbated diastolic dysfunction (DD) induced by hypertension and pressure overload with preserved ejection fraction in female C57BL/6J mice.
2. In post-OVX-treated DD mice, heart weight and fibrosis preceded other metrics of cardiac dysfunction.
3. OVX-induced estrogen-deficiency increased the expression of genes associated with stress, energy metabolism, and especially fibrosis.
4. Metabolic imaging of the heart by positron emission tomography revealed that DD with preserved ejection fraction was associated with mitochondrial dysfunction presenting as increased accretion of cardiac energy substrates, [^18^F]deoxyglucose and [^11^C]palmitate, effects that worsened following OVX.
5. OVX increased cardiomyocyte stiffness and fibrosis and reduced cardiomyocyte lengthening and compliance in the context of female DD.
6. Improved diastolic function following delivery of GPER agonists or estradiol to OVX DD female mice implicates estrogen receptor α in the maintenance of normal cardiac function.

## INTRODUCTION

Heart failure (HF) with preserved ejection fraction (HFpEF) is a common syndrome implicated by high morbidity and mortality and limited treatment options. HFpEF is characterized by diastolic dysfunction (DD) and elevated left ventricular (LV) filling pressure in the presence of HF symptoms in addition to usually elevated natriuretic peptides. While the ejection fraction (EF) remains normal (i.e., typically 50% or greater), patients with HFpEF typically experience DD characterized by ventricular hypertrophy, impaired LV relaxation due to increased cardiac stiffening, and fibrosis. Mechanistically, extracellular fibrosis exacerbates heart stiffening^1^. Low mitral annular velocity (e’), low e’ to late diastolic mitral annular velocity ratio (e’/a’), type 2-4 mitral inflow pattern (E/A), elevated ratio of early mitral inflow velocity E to e’ (E/e’) are echocardiographic Doppler markers for DD HF.

Hypertension, hypertrophic cardiomyopathy, and valvular heart disease increase DD risk^2^. Metabolic comorbidities, i.e., menopause, aging, obesity, and diabetes mellitus^3^, also provoke DD risk. Among women, DD HF prevalence increases during menopause^4,5^, implicating estrogen levels in diastole preservation, since estrogen levels are lower after menopause. We hypothesized that the deficiency in the specific estrogen type, 17β-estradiol (E2), impairs multiple mechanisms and structural features associated with normal diastole. Given that E2 replacement correctly timed with menopause improves diastole, E2-deficiency may also contribute to DD^6,7^. E2, which profoundly impacts metabolism^8^, protects against mitochondrial damage by reducing reactive oxygen species (ROS) production and maintaining oxidative phosphorylation and mitochondrial membrane potential^9^. The active relaxation process of diastole depends on mitochondria-generated ATP^10,11^. Thus, we further hypothesized that E2 counteracts DD by maintaining normal cardiac mitochondrial function.

Elevated angiotensin II (AngII) increases systemic vascular resistance, which forces the already weakened heart to work harder and consume more oxygen^12^ while stimulating fibroblast proliferation and myocyte hypertrophy, manifesting as heart stiffening^13^. Concurrently, increased ROS provokes endothelial dysfunction and further myocardial injury^14,15^. Most mouse models of DD HF feature “two hits,” i.e., stressors of two metabolic pathways. These include obesity-linked insulin resistance plus hypertension^16^, hypertension plus cardiac pressure overload^17^, and obesity-linked insulin resistance plus chronic cardiac pressure overload^18^. Ovariectomy (OVX)-induced E2-deficiency in the HF model comprising obesity-linked insulin resistance plus hypertension failed to increase the severity of multiple HF features.^19^ On the other hand, delivery of a combination of L, HFD, and 4-vinylcyclohexene dioxide, which induces “ovary-intact” menopause to female mice produced a robust DD state.^20^ However, given that ovaries are the major source of testosterone, a concurrent drug-induced androgen excess in this ovary-intact model^21^ was cited as the source of DD induction in this model. Thus, many questions about the effects of E2-deficiency on DD remain unsettled.

Treating male mice with L-arginine analogue N(ω)-nitro-L-arginine methyl ester (L-NAME) plus AngII (L + A) reduces EF to a greater degree than that produced by either agent alone^17^. In contrast, doses of L + A, higher than those used in this study, did not alter EF or fractional shortening (FS) in female mice. Additionally, changes in E/A, E/e′, and e′/a′, characteristic of DD, were more severe in OVX mice treated with L + A with increased diastolic HF severity compared to sham mice treated with L + A due to underlying cardiac mitochondrial dysfunction leading to non-ischemic heart failure in the OVX + (L + A) group^16,17,22^. While OVX in insulin resistant and hypertensive female mice had no effect, we sought to test the hypothesis that OVX in a more severe DD HF model comprising the dual mechanical stressors of L-NAME-induced hypertension and AngII-induced cardiac pressure overload would exacerbate DD and HF.

## METHODS

### Animals and Surgery

Female C57BL/6J mice (15 weeks old) were purchased from Jackson Laboratories (Bar Harbor, ME) and maintained with 12/12 hours of light/dark at room temperature with free access to chow diet (Envigo, Indianapolis, IN, Teklad, 2920x) and water. All studies were approved by the Animal Care and Use Committee of Houston Methodist Research Institute. Mice were randomly assigned to bilateral OVX and sham surgeries, performed as described previously^23^ (**Supplemental Material**). A drop in uterus weight (sham: 0.1 ± 0.006 g, OVX: 0.028 ± 0.005 g, *p* < 0.0001) and levels of plasma E2 measured by ELISA (Calbiotech Co.; sham: 7.628 ± 1.38 pg/mL, OVX: 2.587 ± 0.224 pg/mL, *p* = 0.007, confirmed successful OVX.

### Treatments (See Supplemental Figure S1)

#### L-NAME + AngII to induce DD

Mice were randomly divided into four treatment groups according to sham or OVX surgery and with and without L + A treatment, composed of 0.3 mg/mL L-NAME (Sigma-Aldrich) in 1% saline drinking water and 1.2 mg/kg AngII (Sigma-Aldrich) per day through a subcutaneous osmotic pump^17^ for four weeks. Mouse numbers and data collection points are compiled in **Figure S1A**.

#### Time Course of DD Development

OVX mice were randomly separated into six groups (*n* = 5 per group) and received L + A treatment for either 0, 2, 4, 7, 14, or 21 days. Echocardiography and tissue Doppler analyses were performed at each timepoint (**Figure S1B**).

#### Treatment with Estrogen Receptor-Agonists

OVX + (L + A) mice received the selective estrogen receptor alpha (ERα) agonist propyl pyrazole triyl (PPT, 3.3 µg/day), the selective estrogen receptor beta (ERβ) agonist bis-hydroxyl phenyl propionitrile (DNP, 63 µg/day), the selective G-protein-coupled estrogen receptor (GPER) agonist G-1 (3 µg/day), E2 (4.7 µg/day), or placebo via continuous release subcutaneously pellets (Innovative Research of America, Sarasota, FL) for 21 days (**Figure S1C**).

### Echocardiography

Cardiac function in isoflurane-anesthetized mice was evaluated by transthoracic echocardiography (Fujifilm VisualSonics Inc., Vevo 2100) with a 30-MHz phased array probe and a sweep speed of 100 mm/s^24^. Systolic function was determined from M-mode images in two-dimensional short-axis view at the papillary muscle level. Diastolic function was measured by transmitral pulsed-wave Doppler and tissue Doppler in apical four chamber views. LV longitudinal strain and strain rates were assessed by speckle-tracking echocardiography in two-dimension longitudinal-axis views. At least three continuous recordings on each mouse were manually analyzed independently by two people using Vevo Lab1.7.1 software.

### Mitochondrial Isolation and Respiration

Mitochondria were isolated from freshly excised hearts by differential centrifugation^17^. Mitochondrial function was assessed using a Seahorse XF24 Extracellular Flux Analyzer (Agilent Technologies, Inc., North Billerica, MA) in response to pyruvate-malate (Pyr+Mal, Pyr: 5.0 mM, Mal: 2.0 mM), ADP (1.0 mM), and oligomycin (1.0 µM) using the Cell and Mito-stress kit (Agilent Technologies). Oxygen consumption rates (OCR) in pmol O_2_/min were determined in triplicate and normalized to mitochondrial protein concentration. Mitochondrial morphology (1000×) was determined by transmission electronic microscopy (TEM).

### CM Contractility and Elasticity

CMs were isolated using a Langendorff perfusion system^25^. Briefly, mice were anticoagulated with heparin (1000 IU/kg, intraperitoneal) and euthanized. The heart was rapidly excised, cannulated via the aorta, and mounted onto a Langendorff apparatus (Hugo Sachs Elektronik, Germany). The coronaries were first cleared with a Ca^2+^-free buffer, followed by an enzymatic digestion using a specialized buffer containing type IV collagenase (200 U/mL) at a flow rate of 2.5 mL/min. Once flaccid, the heart was mechanically dissociated to separate the CMs. The cells were filtered and then subjected to a stepwise calcium reintroduction protocol (250 µM–900 µM CaCl_2_) to restore physiological levels. CM pellets were suspended in a 1.8 mM CaCl_2_ buffer for subsequent contraction studies and treatments, including in vitro oligomycin exposure.

Contractility was recorded using IonOptix imaging system (IonOptix Inc., Westwood, MA) in CMs in 1.8mM CaCl_2_ buffer and paced by field-stimulation (10 V, 1.0 Hz)^26^. Myocyte contraction rates (shortening) and relaxation (re-lengthening), analyzed using IonWizard 6.0 software, were expressed as dL/dt. Single exponential analysis of the re-lengthening phase duration was expressed using tau factors. CM elasticity was evaluated by atomic force microscopy (AFM; BioScope Catalyst, Bruker, Santa Barbara, CA)^27^. The force curves from individual myocytes were obtained by the AFM contact probe at the center of the longitudinal axis of the CM. Young’s modulus (kPa) was calculated from more than 100 force curves acquired upon contact with individual myocytes using Nanoscope software (v1.5). Sarcomere lengths were aligned and averaged (IonWizard 6.3; IonOptix, Milton, MA, USA), and the first derivatives were calculated in Excel.

### Positron Emission Tomography Imaging

[^18^F]2-fluoro-2-deoxy-d-glucose ([^18^F]FDG) or [^11^C]palmitate were produced in an on-site cyclotron. After a four-hour fast, mice were anesthetized and injected either intraperitoneally with [^18^F]FDG (∼200 μCi/mouse) or via tail vein with [^11^C]palmitate (∼200 μCi/mouse). Positron emission tomography (PET; 1 hour) and computed tomography (CT, 3 mins) scans were collected immediately using Siemens Inveon PET/CT system (Siemens, Erlangen Germany) and viewed using Inveon Research Workplace software (Siemens). Heart standardized-uptake-values (SUV) were calculated from the magnitude of radio-decay normalized to body weight^28-30^.

### Statistical Analysis

GraphPad Prism 5.0 software (GraphPad Software, San Diego, CA) was used for all statistical analyses and graphic display. Data are expressed as mean ± SEM. One-way ANOVA followed by post-hoc Tukey’s comparison for multiple groups was performed, with *p* < 0.05 considered a significant difference. The number of mice (*n*) is indicated in figure legends.

## RESULTS

### OVX Exacerbates (L + A)-Induced DD with Preserved EF

EF and FS were similar across sham, OVX, (L + A), and OVX + (L + A) groups (**Figure 1A, B**). Stroke volume (SV) and cardiac output, however, were reduced across both (L + A)-treated groups, though most profoundly in the OVX mice (**Figure 1C, D)**. Doppler studies revealed that the ratio E/A, an early DD index, decreased in sham + (L + A) mice but returned to a “normal” range in OVX + (L + A) (**Figure 1E, S3**). The ratio E/e’, representing LV filling pressure, was increased by (L + A) treatment (**Figure 1F**) and increased further by OVX. Although the ratio e’/a’ was not affected by OVX alone, it was reduced by L + A and even more so by the combination OVX and L + A treatment (**Figure 1G**). While treatments had little effect on systole (**Table S1**), the OVX + (L + A) treatment impaired diastole. Further, while CM size was increased by (L + A) treatment, OVX + (L + A) treatment increased CM size to the greatest extent (**Figure 1I, J**). The magnitude of cardiac fibrosis in the OVX + (L + A)-treated group exceeded that observed in the other three groups (**Figure 1H, K**), and when normalized to tibia length, OVX + (L + A) mice vs. the other three groups showed the greatest heart weight, a marker of left ventricular hypertrophy (**Figure 1L, Table S1**). Systolic and diastolic blood pressure were not different for OVX vs. Sham or OVX + (L + A) vs. (L + A) treatments (**Figure 1M, N**). Collectively, these data established that OVX exacerbates multiple biomarkers of DD among (L + A)-treated female mice.

**Figure 1.**
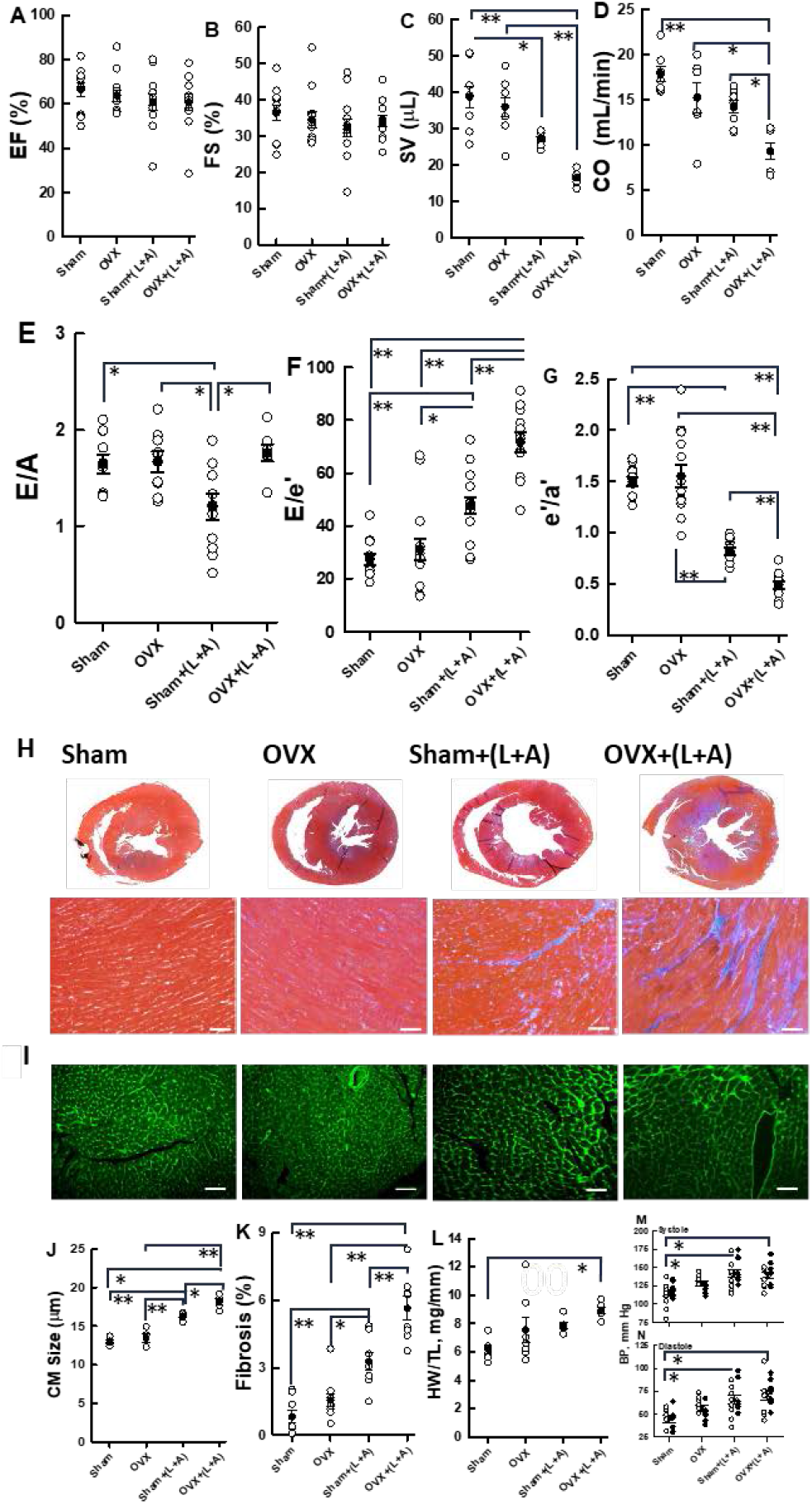
DD markers in ovariectomized mice receiving L and A. **A**, Left ventricular (LV) ejection fraction (EF); **B**, LV fractional shortening % (FS); **C**, stroke volume (SV); **D**, cardiac output (CO); **E**, E/A; **F**, E/e’; **G**, e’/a’; Representative images of H, Masson’s trichrome fibrosis staining; I, WGA staining of LV cross-sections and **J**, CM size, **K**, %fibrosis; **L**, Heart weight. **M**, Systolic and **N**, diastolic blood pressure. Data are presented as the mean ± SEM from 6–8 mice/group; \**p* < 0.05, ** *p* < 0.01.

### OVX + (L + A) Changes Cardiac Structure and Function

The onset and kinetics of cardiac function were determined using serial echocardiographs over 21 days in OVX + (L + A) mice. For comparison, data were normalized to the first time point (**Figure 2;** unnormalized data are provided in **Figure S2)**. While EF and FS remained unaltered (**Figure 2A, B**), we observed an exponential linear decay in SV and CO, both with halftimes of 3.6 days and linear declines and increases respectively in e’/a’ (**Figure 2C, E**) and E/e’ (**Figure 2F**) over 21 days post-treatment. According to the slopes of the lines, rate of decline in e’/a is greater than the rate of increase in E/e.’ According to the fitted curves, the changes in at 2 days for EF, FS, SV, CO, e’/a, and E/e’ were - 1, -1, -10, -12, -13, and +22% (Figure A-F). The linearity of SV and CO and e’/a and E/e’ suggests each dependent variable correlates linearly with another other as illustrated between E/e’ and e’/a’ in **Figure S4**.

**Figure 2.**
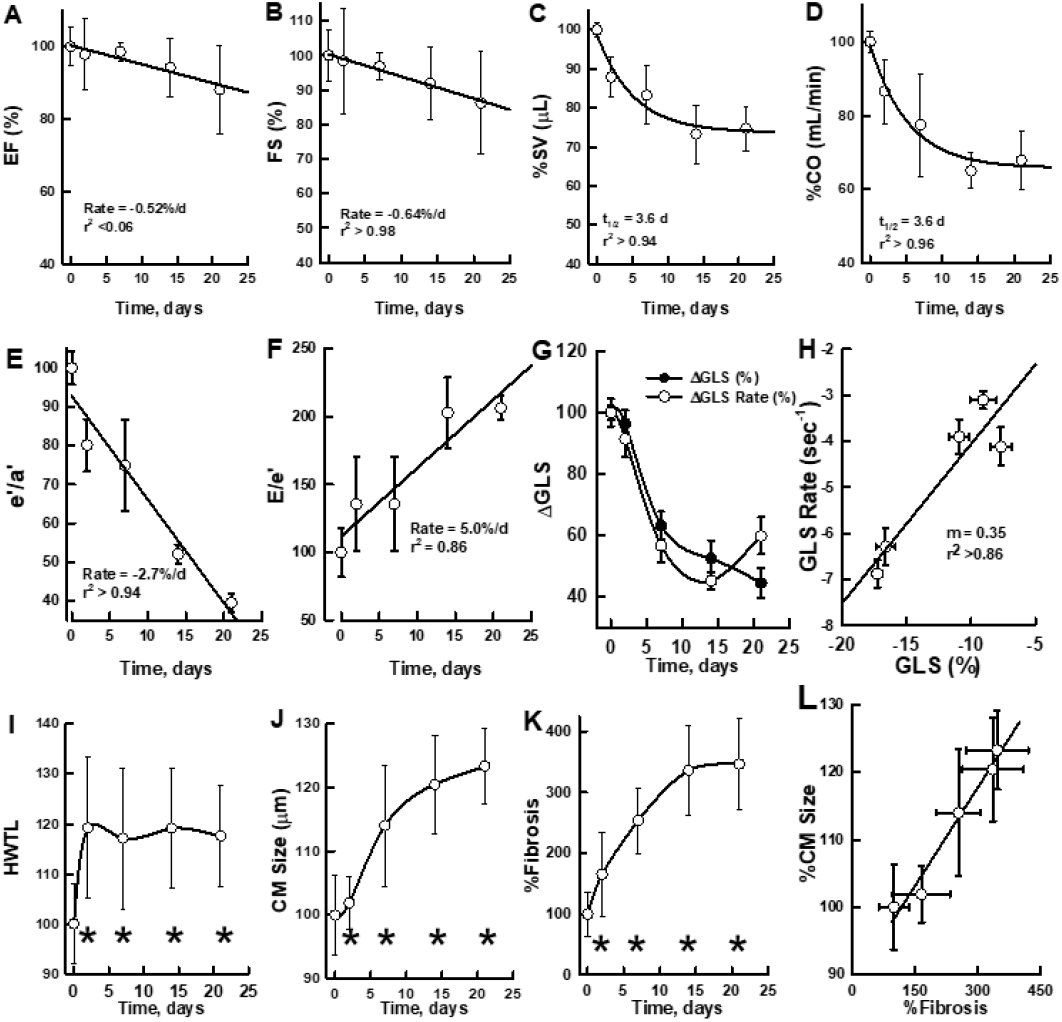
DD markers in OVX + (L + A) mice over time. **A**, LV ejection fraction (EF); **B**, LV fractional shortening (FS), **C**, stroke volume (SV), **D**, cardiac output (CO); **E**, e’/a’; **F**, E/e’; **G**, LV longitudinal strain and strain rates; **H**, Correlation of longitudinal strain vs. strain rates; **I**, heart weight, **J**, CM size, and **K**, fibrosis %, **L**, Correlation of % CM size and % Fibrosis. Data are the mean ± SEM from 6–8 mice/group; \**p* < 0.05 vs. Day-0. Data are normalized to Day 0 = 100%.

OVX + (L + A) treatment induced near-parallel decreases in longitudinal strain and strain rate by day 7, after which both variables plateaued (**Figure 2H**). Hypertrophy appeared on Day 2 and plateaued by Day 21 (**Figure 2I**). While CM size increased on Day 7 (**Figure 2J**), we observed a meaningful increase in fibrosis as early as Day 2, which continued to increase until day 14 (**Figure 2K**). Notably, CM size and level of fibrosis were linearly correlated (**Figure 2L**). Collectively, changes in cardiac function, which began on day 7, became more profound by day 14 post treatment (**Figure 2C–G**); nonlinear functional and structural changes (**Figure 2I–K**) occurred as early as 2 Days post-OVX + (L + A) treatment.

### OVX Amplifies Cardiac Mitochondrial Dysfunction in (L + A)-Treated Mice

Cardiac mitochondrial function in response to pyruvate-malate with ADP revealed that mitochondrial OCR had the largest reduction in the OVX + (L + A) group (**Figure 3A)**. Furthermore, OVX + (L + A) compared to sham mice experienced the greatest increase in mitochondrial ROS and a decrease in mitochondrial DNA content, a metric for cardiac mitochondria number, (**Figure 3B, C)**. Transmission electron microscopy (TEM) of heart tissue from OVX + (L + A)-treated mice revealed rounded and dense mitochondria, arrayed in disorganized networks that contained distinct lipid droplet-like structures (**Figure 3F**). The number of abnormal mitochondria (**Figure 3D)** was significantly higher in OVX + (L + A)-treated mice compared to sham. Finally, in the OVX + (L + A) group, ROS production increased by Day 2 and continued increasing until plateauing on Day 10. Overall, OVX amplifies the mitochondrial morphological and functional abnormalities induced by (L + A) treatment.

**Figure 3.**
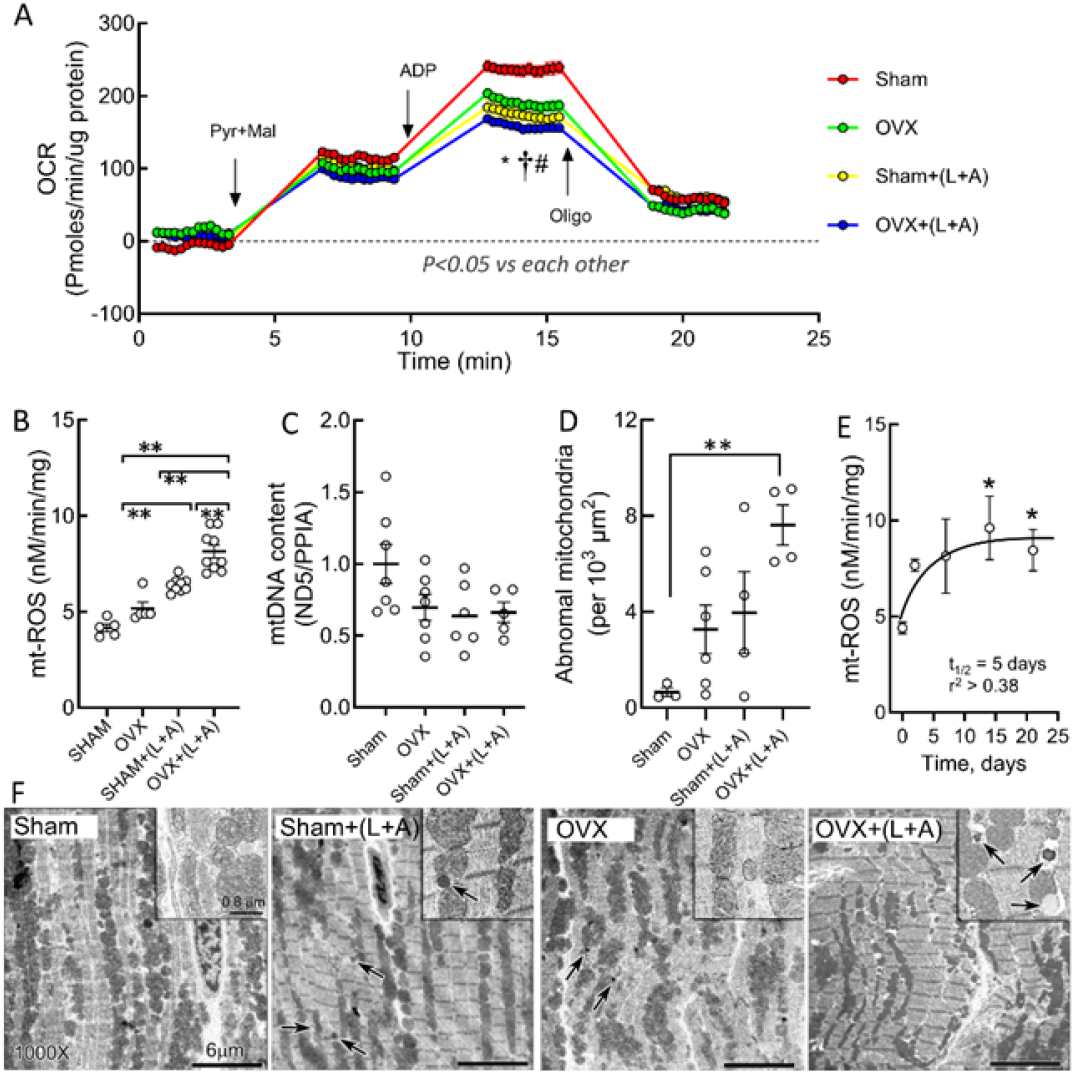
Impaired CM mitochondrial function in DD. **A**, Oxygen consumption rate (OCR) with sequential addition of sodium pyruvate (Pyr, 5 mM) + L-malate (Mal, 2 mM), ADP (1 mM), and oligomycin (Oligo, 1 µM), \**p* < 0.05 vs. sham; #*p* < 0.05 vs. OVX; †*p* < 0.05 vs. sham + (L + A); **B**, ROS in mitochondrial isolates; **C**, mitochondrial DNA (mtDNA) content; **D**, Number of abnormal mitochondria (See F); **E**, Time-dependent changes in ROS with P+M in OVX + (L + A); **F**, Mitochondrial morphology by TEM (Insert, higher magnification; arrows show damaged mitochondria). Data are presented as mean ± SEM, 4–8 mice/group. \*\**p* < 0.01 for **B, D**.

### OVX Increases Expression of Genes of Stress, Energy Substrate Metabolism, and Fibrosis

OVX mice treatment with (L + A) induced significant and meaningful changes in gene expression compared to either OVX or (L + A) treatment alone, which produced mostly small changes in gene expression. OVX + (L + A) treatment increased expression of heart failure genes, including atrial natriuretic peptide, brain natriuretic peptide, and Growth Differentiation Factor 15, which is associated with DD (**Figure 4**)^31^. In contrast, OVX + (L + A) treatment suppressed the expression of mitochondrial biogenesis and function transcription factor Nuclear respiratory factor 1 expression, while OVX alone modestly increased Nuclear respiratory factor 1 expression. Lower peroxisome proliferator activated receptor gamma coactivator 1 alpha expression was observed in both (L + A)-treated groups. Interestingly, OVX alone elicited the opposite effect. OVX + (L + A) also increased the expression of mitochondrial uncoupling protein 3, a protein that responds to oxidative stress as well as the expression of glucose metabolism genes pyruvate dehydrogenase B and lactate dehydrogenase A. Both OVX and OVX + (L + A) treatment increased non-esterified fatty acid (NEFA) uptake and metabolism genes, carnitine palmitoyltransferase 1B, and CD36. OVX + (L + A) also increased expression of tumor necrosis factor alpha, a known inducer of fibrogenesis and gene for the inflammation-associated inflammatory protein^32^; further, these mice responded with a profound increase in expression of fibrosis-associated genes, collagen type 1 α 1 (Col1a1) and collagen type III α 1 (Col3a1), which likely underlies the significant increase in fibrosis. Considered together (**Figure 4**), genes associated with stress response, energy metabolism, and fibrosis are primary targets implicated by hypertensive, pressure-overloaded OVX female mice. The gene expression profiles support a model of stress- and fibrosis-induced severe cardiac dysfunction in OVX + (L + A)-treated mice.

**Figure 4.**
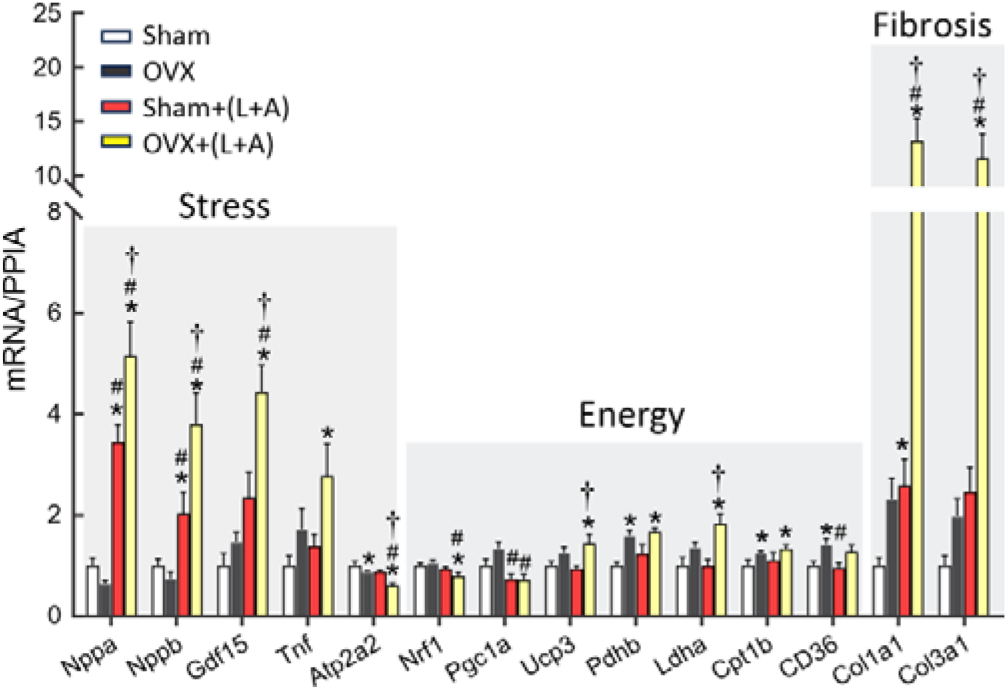
Gene expression OVX+(L+A) mice, as labeled. Relative gene expression, normalized to PPIA. Data presented as mean ± SEM from 6-8 mice/group. *p<0.05 vs. Sham; #p<0.05 vs. OVX; †p<0.05 vs. Sham+(L+A).

### E2-Deficiency Increases In Vivo Cardiac Substrate Accretion

Axial views of PET and PET with CT reveal the differential accumulation of [^11^C]palmitate and [^18^F]FDG in mice across the sham, sham + (L + A), and OVX + (L + A) groups (**Figure 5A**). Compared to sham, cardiac accumulation of [^11^C]palmitate increased by 27% in sham + (L + A) mice and 63% in OVX + (L + A) mice at 10 min after tail vein injection (**Figure 5B**). Cardiac accumulation of [^18^F]FDG increased by 67% in sham + (L + A) mice and 130% in OVX + (L + A) mice at 1 hour after intraperitoneal injection (**Figure 5C**). The increased energy accretion as NEFA and glucose, among OVX + (L + A)-treated mice complements the increased expression of energy-associated genes. However, we were unable to measure rates of energy substrate turnover with either PET tracer.

**Figure 5.**
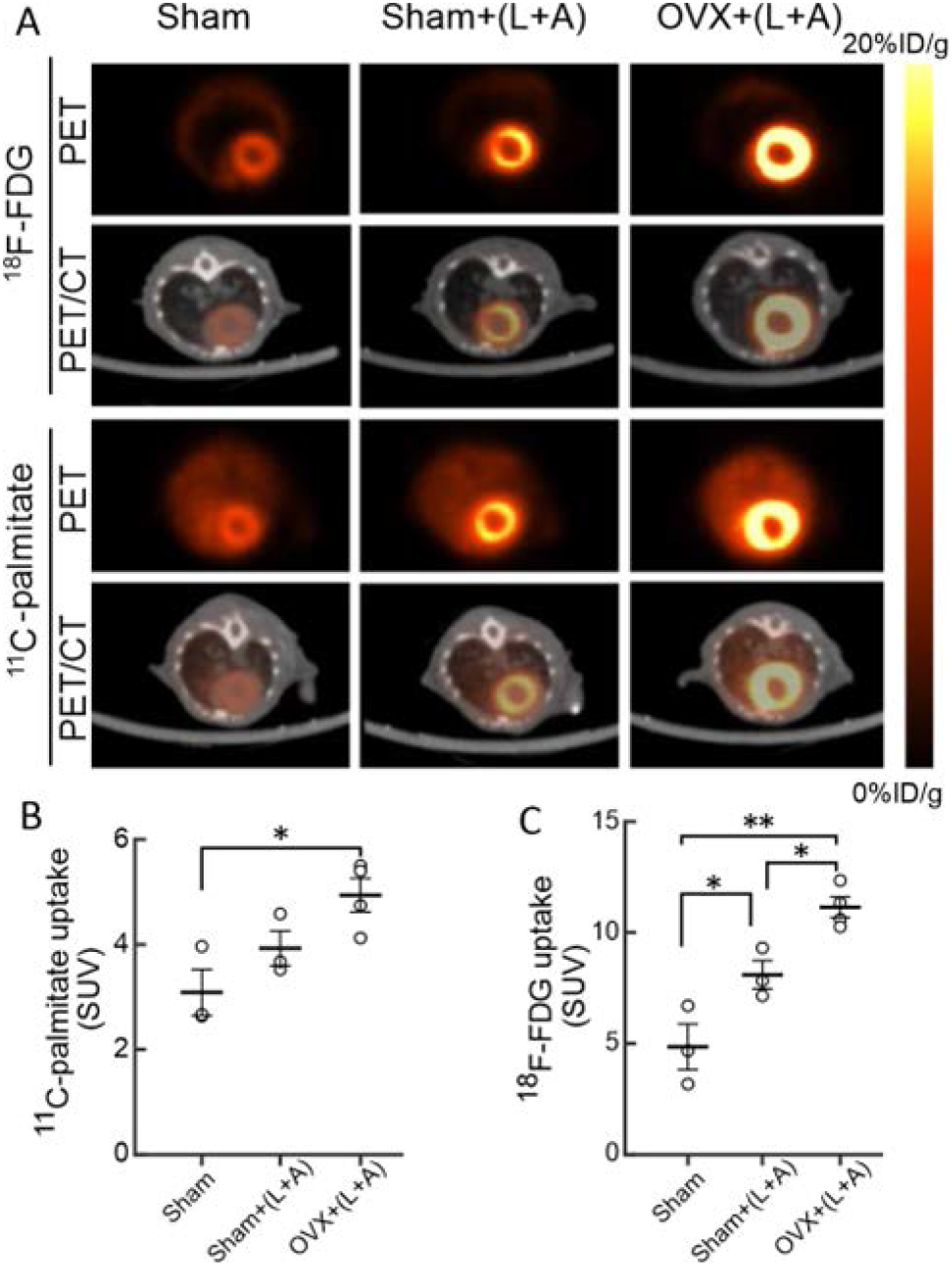
Cardiac Energetics in Mice. **A**, Representative images of axial views of [^18^F]fluorodeoxyglucose ([^18^F]FDG) and [^11^C]palmitate accretion in mice according to positron emission tomography (PET) imaging and computed tomography (CT) as labeled. **B**, Calculated standardized u5ptake values (SUV) for [^11^C]palmitate and **C**, [^18^F]FDG. Data are presented as mean ± SEM from 3–4 mice/group in **B** and **C**. \**p* < 0.05, \*\**p* < 0.01.

### CM Structural Biophysics

We observed that (L+A) treatment profoundly altered CM biophysics. The Young’s modulus, a metric for cardiac stiffening, was increased in both sham + (L + A) and OVX + (L + A) CMs (**Figure 6A**). While CM contraction was not affected (**Figure 6B**) by treatment, both (L + A)-treated groups exhibited decreased CM relaxation (**Figure 6C**) and an increased Tau factor (**Fig 6D**). OVX treatment alone indicated greater cardiac stiffening and CM re-lengthening. We observed no differences in calcium transient assessments, including Ca^2+^ release rate, Ca^2+^ release time, peak, time to peak, decay time, or decay rate among treatment groups (**Table S2**).

**Figure 6.**
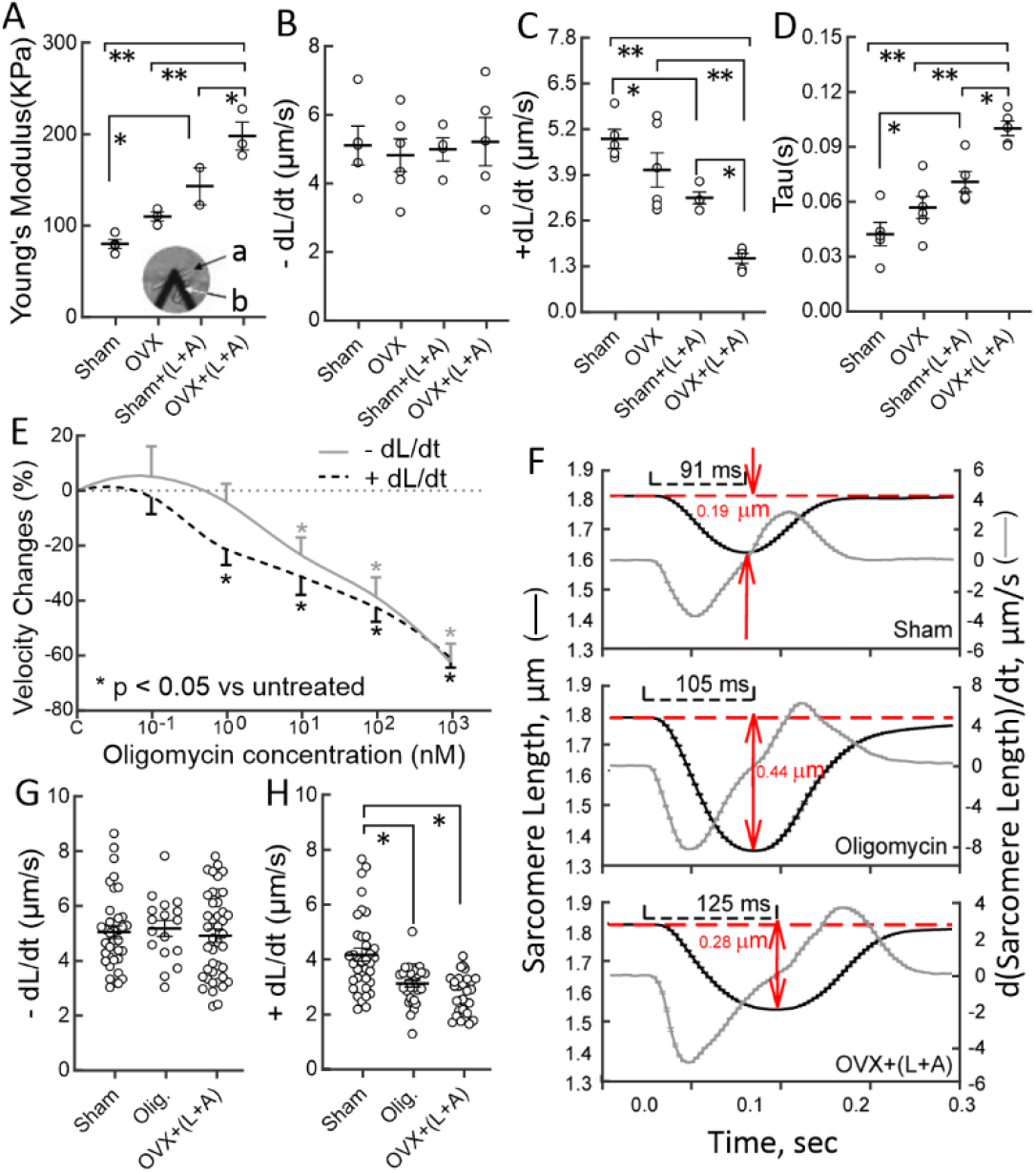
Effects of DD on cardiomyocyte contractility and stiffness. A, Cardiomyocyte (CM) elasticity measured by AFM (AFM Image: a is the CM surface and b is the cantilever); B, CM contraction (-dL/dt), C, CM re-lengthening (+dL/dt, relaxation), and D; Tau factor; E, Dose-response of oligomycin to CM contractility; F’ Mean sarcomere length over time and its first derivative (ds/dt = 0, gray lines); G, CM contraction (-dL/dt), and H, CM relaxation (+dL/dt) in sham, oligo-treated, and OVX + (L + A) groups. Data presented as mean ± SEM from 4–5 mice/group. *p < 0.05; **p < 0.01.

Oligomycin treatment of CMs isolated from the sham group reduced both contraction and relaxation with the effects on relaxation appearing at a lower oligomycin concentration (∼1 nM), and contraction only affected at ≥10 nM oligomycin (**Figure 6E**). Compared to sham-treated mice, sarcomere rates of change in CM length were similarly reduced in the OVX + (L + A)-treated mice and in mice in which ATP synthesis was inhibited by oligomycin (**Figure 6G, H**). According to the derivative of sarcomere length with respect to time, the times to minimum CM length increased as sham < oligomycin < OVX + (L + A), respectively, as 0.091 s, 0.105 s, and 0.125 s, while minimum lengths were 0.19, 0.44, and 0.28 µm (**Figure 6F**). Collectively, stiffened CMs and delayed re-lengthening due to impaired mitochondrial ATP synthesis may be among the key cellular mechanisms underlying DD.

### Role of Estrogen Receptors in Diastolic Function

We determined the effects of Erα (PPT), Erβ (DPN), and GPER (G-1) agonists as well as E2 delivered subcutaneously to OVX + (L + A)-treated mice on cardiac function for 21 days (**Figure S1C**). All treatments except DPN suppressed the OVX + (L + A)-induced alterations of cardiac function. E2, PPT, and G-1 did not affect EF (**Fig. 7A**) but increased SV (**Figure 7B**) and cardiac output (**Figure 7C**). Similarly, E2, PPT, and G-1 treatments improved both E/A (**Figure 7D**) and e’/a’ (**Figure 7F**) and reduced E/e’ (**Figure 7E**). Systolic blood pressure was lower for PPT and DPN compared to placebo, and diastolic blood pressure was lower for PPT compared to placebo (**Figure 7G, H**). Collectively, these data implicate ERα and GPER in the modulation of E2 depletion-dependent DD.

**Figure 7.**
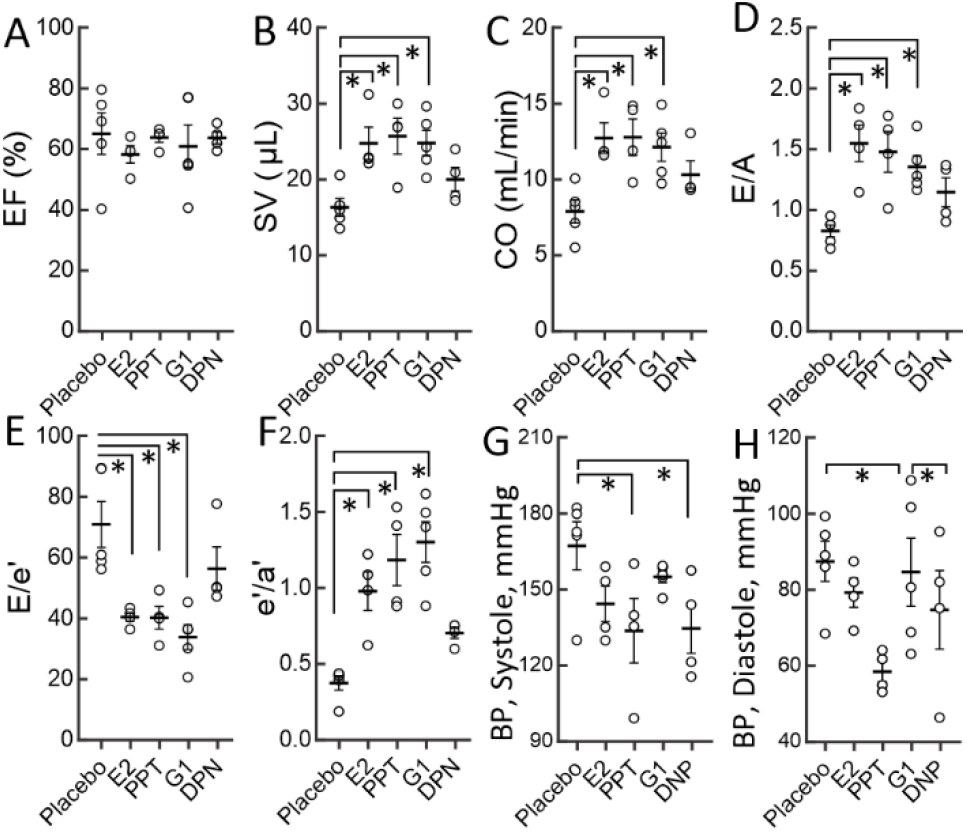
Effects of Estrogen Receptors on Cardiac Function in OVX + (L + A) Mice. **A**, EF, **B**, SV, **C**, CO, **D**, E/A, **E**, E/e’, **F**, e’/a’, **G**, systolic BP (blood pressure), and **H**, diastolic BP were measured in (L + A)-treated OVX mice treated with E2, PPT, G-1, DPN, and placebo. Data are presented as mean ± SEM from 4–5 mice/group, \**p* < 0.05.

## DISCUSSION

The present findings complement our earlier observations that selective pharmacological ERα activation in OVX female Ldlr^-/-^ mice fed a high-fat diet (HFD) suppresses weight gain, improves insulin sensitivity and cardiac mitochondrial function, reduces visceral fat accumulation, and induces expression of metabolic genes in the heart^9^. The dual insults of a HFD and hypertension induced by inhibiting constitutive nitric oxide synthase with L-NAME in mice, models multiple features of human DD HF. However, OVX-induced E2-deficiency in female mice receiving HFD and L-NAME does not alter multiple DD HF markers^16,19^. In male mice, the dual insults of (L + A), which respectively induce hypertension and cardiac pressure overload, are associated with impaired mitochondrial respiratory function and increased oxidative stress compared with either agent alone. This treatment combination results in more severe, nonischemic HF than that induced by the combination of L and HFD. Mouse models of DD HF have been critically assessed,^33^ including the multi-hit mouse models of (HFD + L) and (HFD + A), which share a phenotype of obesity and insulin resistance. Our model, (L + A), recapitulates the DD HF phenotype without obesity or insulin resistance. By one measure of HF, E/e’, the effects of (L + A) were more severe than those reported for (HFD + L)^16,19^ and (HFD + A)^34^ models.

Using multiple approaches, we tested the effects of OVX on multiple markers of DD HF in a mouse model induced by (L + A). While having no effect on EF and SV (**Figure 1A, B**), markers of DD, i.e., SV, CO, E/e’ and e’/a’, increased in severity from sham to OVX alone to sham + (L + A) to OVX + (L + A) (**Figure 1C–L**). Although the E/A ratio appeared in the nominally ‘normal’ to slightly elevated range in the OVX + (L + A) group, this may be the deceptive consequences of a “pseudo-normalization” based on the increased ratio of E/e’ (**Figure 1E**)^10,35^.

Changes in DD markers following each treatment were also time dependent. The halftimes for changes in SV and CO were similar and given that SV is a determinant of CO, this would be expected. The linear changes in e’/a’ and E/e’ over 21 days positively and negatively correlated with time, as expected given that e’ is in the numerator and denominator of the respective dependent variables (**Figure 2E, F**), e’/a’ and E/e’ correlated (**Figure S4**), suggesting a common underlying cause. Other changes in DD metrics were nonlinear with respect to time. Heart weight plateaued at a 20% increase by Day 2 post treatment, and despite their nonlinearity with time, longitudinal strain and strain rates positively correlated (**Figure 2H**), suggesting a common etiology originating in fibrosis. Fibrosis appeared earlier and with greater severity (+65% by day 2 post treatment) than other DD markers and increased nonlinearly reaching 250% by day 21 (**Figure 1K, 2K**).

Changes in mitochondrial function followed a similar pattern of increasing dysfunction with treatment time and severity. OCR, ROS, and the number of abnormal mitochondria increased while mitochondrial DNA content declined. Relative to sham treatment, cardiac accumulation of [^11^C]palmitate increased by 27% and 63% following sham + (L + A) and OVX + (L + A), respectively. Cardiac accumulation of [^18^F]FDG increased by 67% and 130% (**Figure 5**), due in part to reduced energy utilization as revealed by a reduced OCR. The increased *in vivo* energy accretion via NEFA and glucose and the reduced oxygen consumption signifies a severe metabolic imbalance. This may reflect a transfer from an adult cardiac gene program that depends on the NEFA utilization to a more oxygen-efficient fetal glucose-dependent gene program of a stressed heart^28^. This could also be mechanistically linked to the parallel changes in OCR (**Figure 3A**), which also declined most substantially in the OVX + (L + A) group. While our study did not assess the downstream utilization of the substrates, the observation that both NEFA and glucose uptake increased during mitochondrial insufficiency and impaired cardiac function suggests myocardial ATP-depletion despite excess substrate availability in DD. Parallel changes were observed in other markers of mitochondrial failure that may impair energy utilization, including declining mitochondrial DNA content, the appearance of abnormal mitochondria, and increased ROS production over time (**Figure 3**).

Fibrosis-associated genes, Col1a1, Col3a1, and Tnf, increased to a greater extent, than stress- and energetics-related genes (**Figure 4**), an observation that comports with the early and severe increase in fibrosis (**Figure 2K**). Although the compensatory increase in CM size was smaller than the change in fibrosis, the two metrics were linearly correlated (**Figure 2L**), consistent with their shared molecular triggers that increase heart mass^36^. While increased heart weight is often driven by the enlargement of individual CMs, fibrosis contributes both directly to total heart mass and indirectly to the mechanical stressors that trigger growth. Further, while myocardial water content is a minor source of the failing heart mass, it contributes disproportionately to impaired cardiac output^14^.

The oligomycin dose-response experiments revealed that CM relaxation is more sensitive to the decline in mitochondrial ATP formation than contraction is, indicating the modest decline in mitochondrial function by OVX + (L + A) treatment is associated with impaired CM relaxation but not contraction. While oligomycin and OVX + (L + A) treatments both reduced the rate of change of sarcomere length, the latter responded the slowest and persisted the longest (**Figure 6D**). As expected, these changes comport with other biometric changes in CM function that appeared more substantially in OVX + (L + A) mice, including increased Young’s modulus (cardiac stiffness), increased tau (enhanced pressure decay during isovolumetric relaxation), and diminished CM lengthening.

DD is also associated with increased mitochondrial ROS production, which triggers inflammation via tumor necrosis factor α and fibrogenesis pathways via tumor growth factor beta and collagen genes Col1a1 and Col3a1^1^. ROS also modulates kinase-phosphatase pathways, which reduce titin compliance^37^ and CM elasticity. Concomitantly, fibrosis in OVX + (L + A) increases LV stiffness and reduces compliance. Inexplicably, there were no differences in calcium reuptake across groups, and given that CM passive stiffness can mediate DD despite improved Ca^2+^ handling, it may not be a critical component of DD. Notably, improved calcium handling produces better outcomes without reducing DD HF symptoms^38^.

Given that OVX treatment alone or in combination with HFD does not provoke DD, a more severe metabolic insult is required to achieve DD. In the current model, these insults are hypertension and pressure overload, which present in older women with DD HF^5,35^. The discordant effects of no difference in DD (E/e’) between sham and OVX mice in their DD HF model^19^ reflect the distinct etiologies of the stressors utilized. While other researchers have employed HFD to simulate metabolic stress, our approach uses Ang II to induce pressure overload. Since the absence of estrogen activates fibrotic genes and enhances CM stiffness following OVX + (L + A) treatment, our data suggests that E2 protects against Ang II-driven pressure overload and supports a model (**Figure 8**) that includes the appearance of a cluster of abnormalities associated with DD in the context of menopause. Data showing the early appearance of fibrosis compared to other HF markers and the largest increase in fibrotic gene expression provoke a mechanistic model that begins with E2-deficiency activating Col1a1 and Col3a1 and in turn, triggering production of collagen type 1α1 and IIIα1 and inducing the known sequala associated with DD-associated DD HF. Expression of tumor necrosis factor (TNF), the master provocateur of inflammation-driven DD in DD HF, could also amplify the phenotype of the OVX + (L + A)-treated mice. TNF’s downstream targets of TGF-β, a driver of collagen synthesis and its disruption of matrix metalloproteinases and inhibitors increase remodeling and CM stiffness and impair CM relaxation. However, the magnitude of TNF’s influence on DD may be overshadowed by the highly expressed fibrotic genes, and thus difficult to estimate.

**Figure 8.**
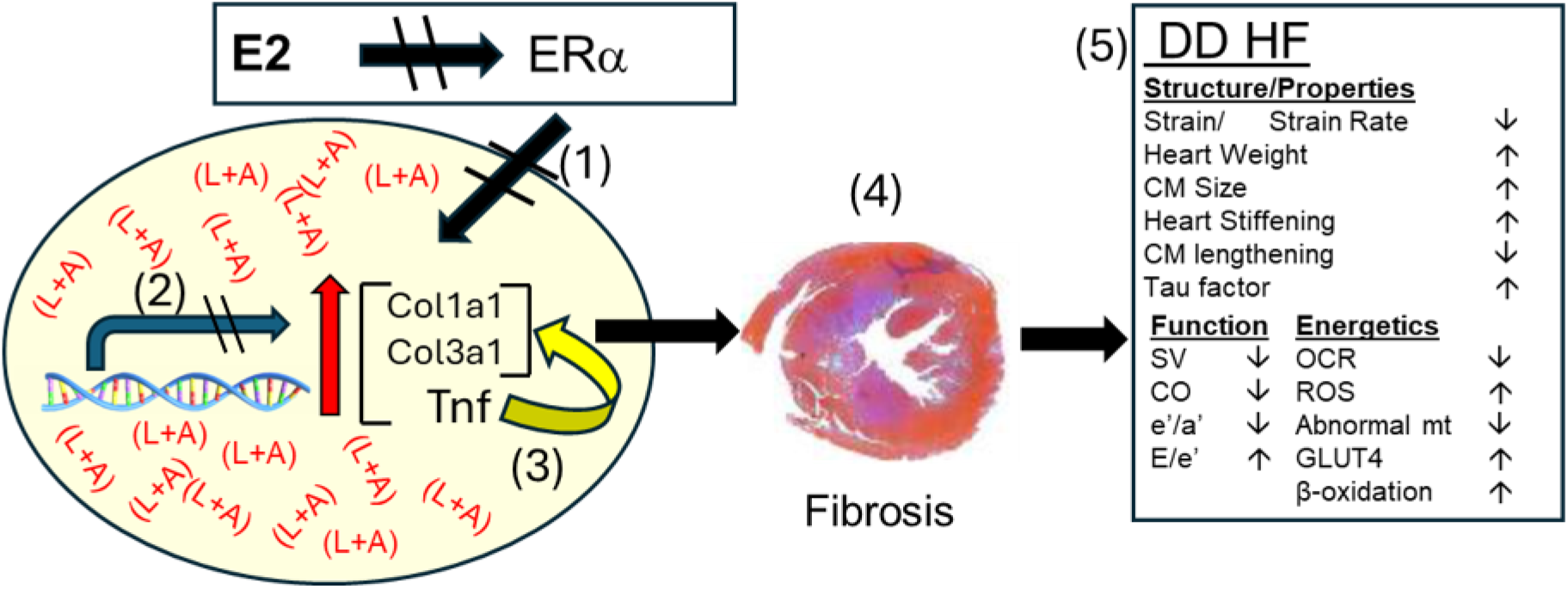
Model of DD-HFpEF with Underlying E2-Deficiency. Without ERα activation (1), fibrotic gene expression is not suppressed and Tnf (2) is activated in turn further activating Col1a1 and Col3a1 (3), which triggers production of collagen type 1α1 and IIIα1 and attendant fibrosis (4). Fibrosis then effects changes in cardiomyocyte structure, properties, and energetics that ultimately impair function and produce the DD-mediated HFpEF phenotype (5).

Menopause-associated E2-deficiency elicits multiple HF comorbidities^8^, including weight gain, insulin resistance, reduced metabolism, and increased cardiovascular disease. The creatine phosphate/ATP ratio is also lower in patients with DD HF^11^. Although our data show functional and structural abnormalities in cardiac mitochondria from OVX + (L + A) mice, these changes are not as extreme as those observed in systolic HF^17^. While the need for mitochondrial ATP for contraction is understood, the need for mitochondrial ATP for diastolic relaxation is not widely acknowledged. ATP powers ion pumps (SERCA2) that transfer Ca^2+^ out of the cytosol during diastole, provide energy for the sarcolemmal Na-K-ATP pump for CM relaxation; its interaction with the actin-myosin bridge dissociates actin and myosin, a step critical to relaxation^10^. Small ATP depletions do not prevent the formation of actin-myosin bridges, and contraction continues due to the high affinity of the myosin head for ATP. However, small reductions in the ATP concentrations reduce actin-myosin dissociation and impair relaxation. Thus, small ATP depletions are known to affect diastole more than systole, whereas larger decreases affect contraction as well as relaxation^10,39^.

Available options for reducing DD HF morbidity center on palliation with loop diuretics, candesartan and spironolactone to reduce blood pressure and block angiotensin and aldosterone effects^40^. The development of more effective DD HF therapeutics has been hindered by the complex multi-factorial pathophysiology, a highly under-investigated gender bias in prevalence, and the dearth of appropriate pre-clinical research models. Our findings provoke future studies that would identify the fate of the substrates within the myocardium and the signaling pathways regulating their utilization. To draw a parallel, increased substrate uptake in skeletal muscle is accompanied by incomplete oxidation, acylcarnitine accretion, mitochondrial uncoupling, and ROS production, thereby inhibiting substrate catabolism^41^. Lur study provides a mechanistic framework for the design of ER-specific, metabolic therapies for DD HF such as sodium-glucose cotransporter 2 inhibitors, which suppress DD in humans with type 2 diabetes ^42^.

## Contributor Roles

1. Conceptualization: AG, KK, HT, DH
2. Data curation: SL, HP, IV, AZ, SC, YW, JG, GR, AG,
3. Formal analysis: AZ, HP, SL, YW, AG
4. Funding acquisition: DH
5. Investigation: AG, KK, DH, IV
6. Methodology: SL, AZ, KK, AG
7. Project direction: AG
8. Resources: GR, AG
9. Software: N/A
10. Supervision: GR, DH, AG
11. Validation: AG
12. Visualization: HP, SL, AZ, AG
13. Writing – original draft: HP, SL, AG
14. Writing – review & editing: HP, SL, IV, AZ, SC, YW, GR, HT, KK, DH, AG

## ACKNOWLEDGEMENTS

The authors thank Houston Methodist Research Pathology, cyclotron, preclinical imaging, AFM cores, and the Comparative Medicine Program for their services. DJH acknowledges Elaine Finger and Marvy Finger for their translational research endowed chair support. HJP acknowledges support from the Dr. Antonio M. Gotto, Jr. Fund. The authors thank Colleen O’Connor, PhD, Office of Faculty and Research Development, Houston Methodist Academic Institute, for scientific editing.

---

**Figure S1.**
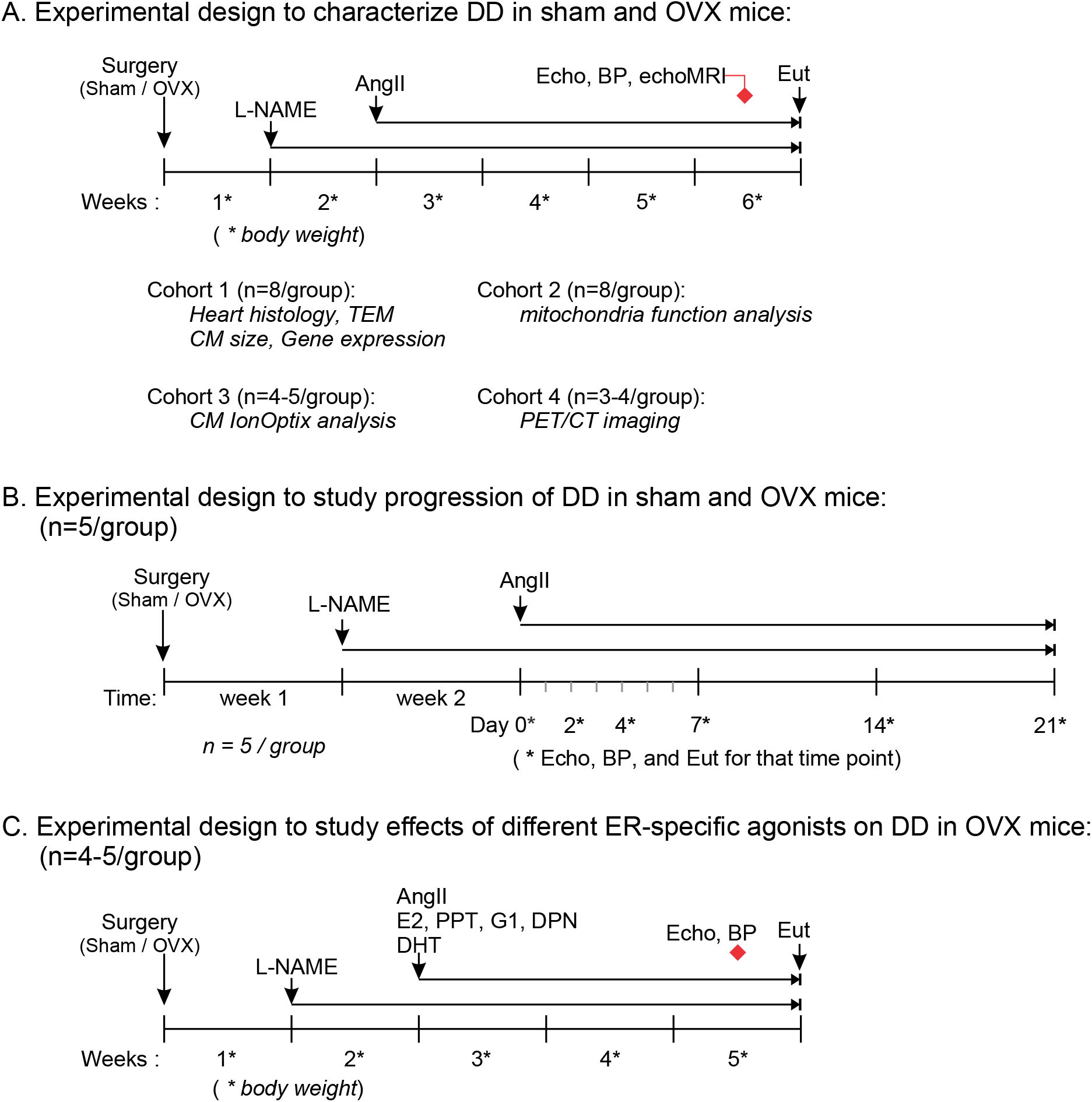
Schematic of Experimental Workflow. Fifteen-week-old female C57BL6 mice underwent ovariectomies or sham surgeries. After one week, half of the OVX mice and half of the sham mice were treated with L-NAME (L, 0.3mg/mL in driving water). A week later, mice received angiotensin II (A, 1.2 mg/kg per day via subcutaneous osmotic pumps). Mice received (L+A) for four more weeks. Four separate experiments were organized: *(A)* Experiment 1 included sham, (L+A) treated sham, OVX, (L+A) treated OVX mice. Four cohorts of mice were required for all assessments listed in the study; *(B)* Experiment 2 was a time-course of (L+A) in OVX mice; *(C)* Experiment 3 was carried out in OVX (L+A) with or without administering either of ER-specific agonists (E2, PPT for ERα, DPN for ERβ, G-1 for GPER), which were delivered subcutaneously in implanted 3-week release pellets. BW: body weight, OVX: ovariectomy, L-NAME: L-*N*G-nitro-arginine methyl ester, Ang II: angiotensin II, Echo: echocardiography, BP: blood pressure; Eut: mice euthanized.

**Figure S2.**
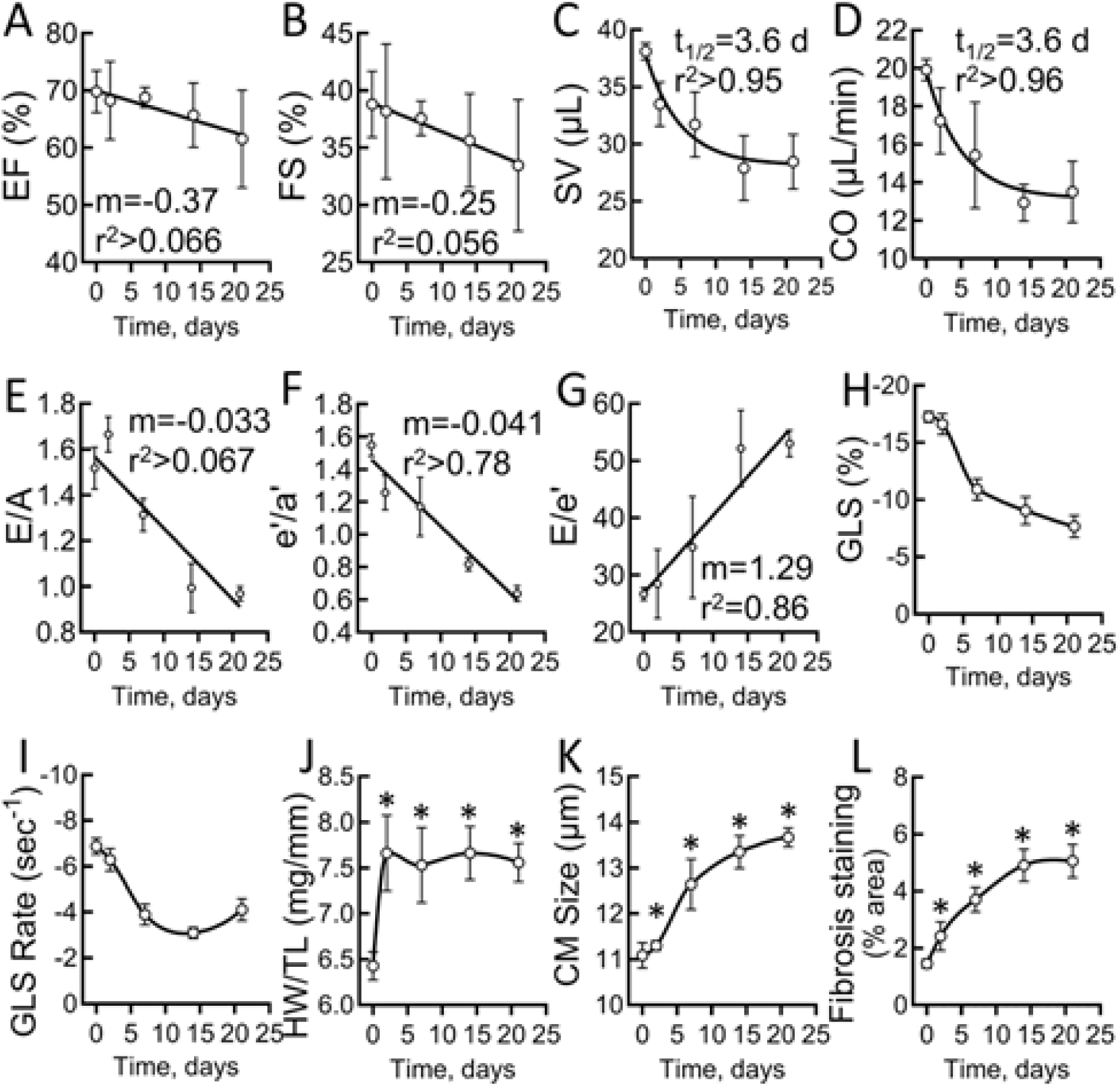
Progression of cardiac impairment in (L+A)-treated OVX mice. *(A)* LV ejection fraction (EF), *(B)* LV fractional shortening (FS), *(C)* stroke volume (SV), *(D)* cardiac output (CO) measured using M-mode echocardiography, *(E)* E/A ratio, *(F)* E/e’ ratio, *(G)* e’/a’ ratio measured by pulsed-wave Doppler and tissue Doppler, *(H)* LV longitudinal strain, *(I)* LV strain rates, *(J)* heart weight to tibia length ratio, *(K)* cardiomyocyte size, and *(L)* fibrosis % are shown. Data presented as mean ± SEM from 5 mice/group. \**p*<0.05 vs. Day-0.

**Figure S3.**
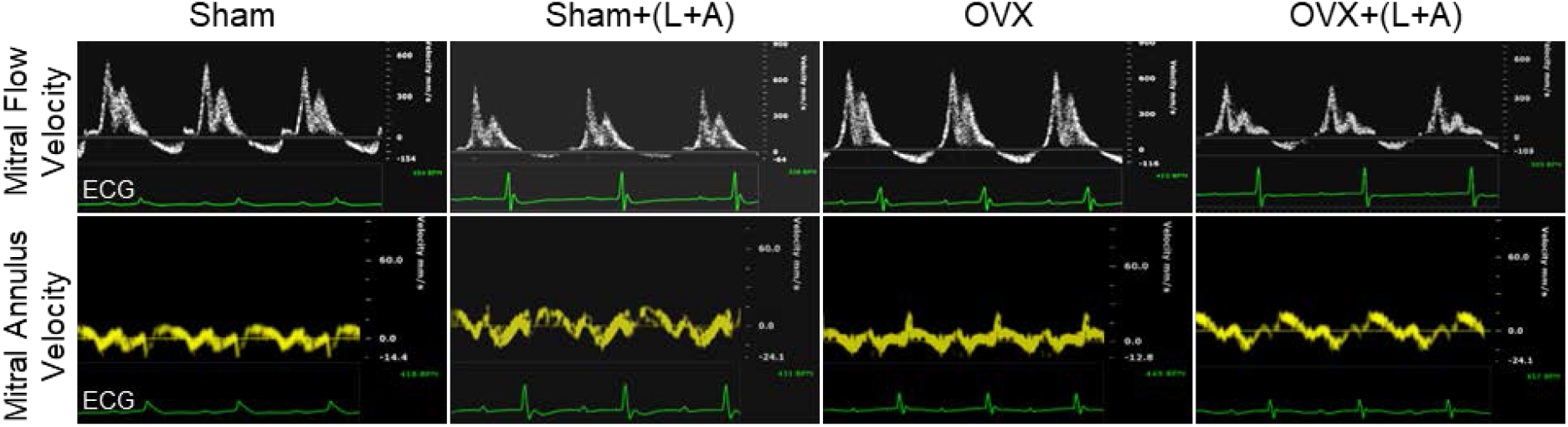
Representative images from pulsed-wave Doppler and tissue Doppler. LV diastolic filling velocities (upper part) were determined by transmitral flow Doppler echocardiography and LV diastolic myocardial velocities (lower part) were measured by tissue Doppler. ECG was shown at the bottom of each Doppler. OVX: ovariectomy; L: L-*N*G-nitro-arginine methyl ester; A: angiotensin II.

**Figure S4.**
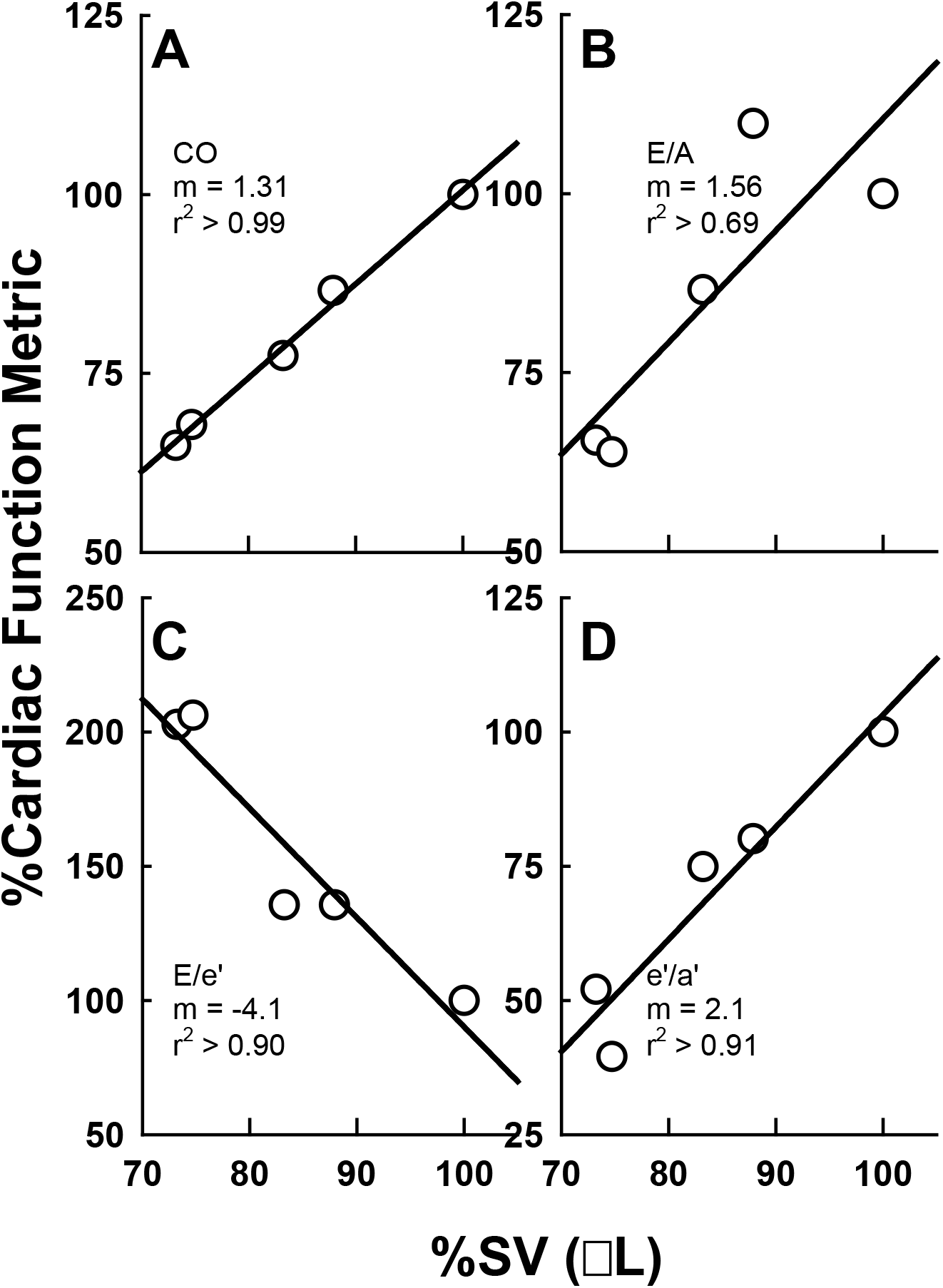
Correlation of Multiple DD Markers with SV.

## SUPPLEMENTAL MATERIAL

**Supplemental Table S1.** Phenotype Summary.

| Parameters | Sham | OVX | Sham+(L+A) | OVX+(L+A) |
| --- | --- | --- | --- | --- |
| Systolic BP (mmHg) | 120.91±3.87 | 119.16±2.52 | 150.51±7.22* | 146.94±5.19* |
| Diastolic BP (mmHg) | 45.63±4.32 | 50.49±3.73 | 71.79±9.7* | 75.31±5.51* |
| Free water (%) | 0.28±0.04 | 0.29±0.05 | 0.28±0.05 | 0.53±0.07*†# |
| LVIDd (mm) | 3.26±0.17 | 3.69±0.18 | 3.64±0.13 | 3.28±0.2 |
| LVIDs (mm) | 2.06±0.13 | 2.49±0.15 | 2.49±0.08* | 2.16±0.15 |
| LVPWd (mm) | 0.94±0.13 | 1.06±0.08 | 0.91±0.08 | 1.32±0.2† |
| LVPWs (mm) | 1.17±0.06 | 1.54±0.07* | 1.14±0.11 | 1.51±0.14† |
| LV Mass corrected(mg) | 80.91±5.53 | 106.59±16.8 | 136.35±25.77* | 126.2±14.51# |
| EDV (uL) | 43.87±4.95 | 58.99±6.61 | 56.69±5.2* | 44.49±6.37†# |
| ESV (uL) | 14.36±2.25 | 22.88±3.22 | 22.34±1.95* | 15.96±2.71† |
| HR (bpm) | 453.71±12.86 | 417.67±18.66 | 438.14±12.25 | 453.6±30.16 |
| MV E (mm/s) | 472.45±19.23 | 455.85±26.58 | 410.15±23.01 | 485.47±27.28 |
| MV A (mm/s) | 291.31±10.59 | 282.81±30 | 374.98±18.18* | 267.76±11.64*# |
| DTE (ms) | 16.93±1.53 | 20.63±1.47 | 22.06±2.92* | 12.17±0.98†# |
| DRE (m/s) | 41.03±12.97 | 41.67±13.89 | 25.1±7.57* | 54.24±20.5†# |
| MV E/A | 1.65±0.09 | 1.67±0.10 | 1.2±0.34* | 1.76±0.10# |
| S' (mm/s) | 19.06±3.32 | 20.34±3.46 | 19.31±3.16 | 21.79±2.24 |
| e' (mm/s) | 19.75±0.54 | 23.36±1.75 | 12.12±4.95* | 7.45±0.5*†# |
| a' (mm/s) | 13.99±0.72 | 14.96±0.46 | 22.66±9.25* | 20.83±1.23*# |

Data are mean ± SEM and are derived from ten to fifteen female mice in each group. BP, blood pressure; % free water by EchoMRI of whole body; LV, left ventricular; LVIDd and LVIDs, LV internal diameter at end of diastole and systole; LVPWd and LVPWs, LV posterior wall thickness at diastole and systole; EDV, end-diastolic volume; ESV, end-systolic volume; HR, heart rate measured under isoflurane-induced anesthesia during echocardiography; MV, mitral valve; E, peak flow velocity of early LV filling; A, peak flow velocity of atrial contraction; DTE or DRE, deceleration time or rate of mitral E wave; E/A, Mitral valve E to A ratio; S', peak velocity of myocardial contraction; e', early peak velocity of myocardial relaxation; a', peak velocity of late atrial contraction. \* $p < 0.05$ vs. Sham. † $p < 0.05$ vs. Sham + (L + A). # $p < 0.05$ vs. OVX.

**Supplemental Table S2.** Summary of Calcium Transient Assessment:

| Parameters | Sham | OVX | Sham+(L+A) | OVX+(L+A) |
| --- | --- | --- | --- | --- |
| Ca<sup>2+</sup> Release Rate (fau/s) | 7,564.38±1,760.80 | 7,314.95±1,093.54 | 8,245.56±1,408.92 | 6,088.42±1,388.07 |
| Ca<sup>2+</sup> Release Time (s) | 0.07±0.00 | 0.04±0.01 | 0.03±0.00 | 0.04±0.01 |
| Peak Ca<sup>2+</sup> (fau) | 170.06±40.51 | 160.04±18.06 | 173.41±48.56 | 120.50±32.60 |
| Time to peak (sec) | 0.08±0.02 | 0.09±0.01 | 0.08±0.01 | 0.10±0.02 |
| Decay Time (sec) | 0.12±0.02 | 0.13±0.01 | 0.13±0.03 | 0.17±0.03 |
| Ca<sup>2+</sup> Decay Rate (fau/s) | 1,007.74±60.01 | 832.94±92.56 | 1,031.82±10.09 | 623.95±147.11 |
| Time to 10% Ca<sup>2+</sup> Rise (sec) | 0.04±0.01 | 0.04±0.00 | 0.03±0.00 | 0.05±0.01 |
| Time to 50% Ca<sup>2+</sup> Rise (sec) | 0.05±0.02 | 0.05±0.01 | 0.05±0.02 | 0.05±0.01 |
| Time to 90% Ca<sup>2+</sup> Rise (sec) | 0.06±0.02 | 0.06±0.01 | 0.06±0.01 | 0.07±0.02 |

Data are mean ± SEM and are derived from three female mice in each group. There were no significant differences between the four groups.

## SUPPLEMENTAL METHODS

OVX and Sham surgeries: Briefly, mice were anesthetized with 2% isoflurane and a skin incision (∼2 cm) was made at the dorsal midline after sterilization. Ovaries were located and removed by cauterization (Bovie Medical Corp., FL). The incisions in muscle and skin were closed sequentially with an absorptive suture (Ethicon, Inc.). Sham-operated mice were subjected to the same procedure except for the excision of ovaries. All surgical procedures were performed under aseptic conditions. Carprofen (0.1mg/kg, subcutaneous), buprenorphine SR (0.6mg/kg, subcutaneous), and bupivacaine (10µl at local incision) were administrated to manage post-operative pain. Body weights and body composition (EchoMRI LLC, Houston, TX) was assessed periodically to monitor wellness of the mice.

Blood pressure was assessed by non-invasive tail-cuff method using BP-2000 Blood Pressure Analysis System (Visitech Systems, Inc. Apex, NC).

EchoMRI: Body composition was analyzed by EchoMRI system (EchoMRI LLC, Houston, TX) and free water indicative of passive congestion was expressed as a percentage of body weight.

Tissue collection: Mice were sacrificed by cervical dislocation and the excised heart was sectioned into 3 pieces, and either fixed in 10% neutral buffered formalin, or snap frozen into liquid nitrogen or stored into biological preservative solution (BIOPS, 2.77 mM CaK_2_EGTA, 7.33 mM K_2_EGTA, 5.77mM Na_2_ATP, 6.56mM MgCl_2_·6H_2_O, 20mM Taurine, 15mM Na_2_Phosphocreatine, 20mM imidazole, 0.5mM dithiothreitol, and 50mM MES, pH7.1) for isolation of cardiac mitochondria. Another batch of tissue samples were fixed for transmission electronic microscopy (TEM) in 2.5% glutaraldehyde solution and processed for examination of mitochondrial morphology.

Assessment of cardiac fibrosis and myocyte size: Paraffin-embedded heart tissue were cut into 4.0 µM sections and stained with Masson’s Trichrome staining kit (Sigma-Aldrich, St. Louis, MO)^1^ for cardiac fibrosis and stained with wheat germ agglutinin (WGA)-FITC conjugate staining kit (Sigma-Aldrich, St. Louis, MO)^2^ to quantitate myocyte size. The area of fibrosis tissue was analyzed by Image J and expressed by areas % of fibrotic tissue over total tissue per sample in images captured by bright-field microscopy (Nikon, Inc. Tokyo, Japan). Images of the WGA-stained heart sections were captured by fluorescence microscopy (Nikon, Inc. Tokyo, Japan) with 488/510 of excitation/emission and analyzed by NIS elements software (Nikon, Inc. Tokyo, Japan). At least 300 cells were manually analyzed from 3 to 5 fields per sample by measuring shortest diameters. The averaged diameter was used for each sample expressed in µm.

Mitochondrial Isolation: Briefly, fresh heart tissue was washed, minced and then homogenized in buffer A (220 mM mannitol, 70 mM sucrose, 5mM MOPS, pH 7.4). The homogenate was centrifuged at 800g for 10 min at 4°C in buffer B (2mM EGTA and 5% fatty acid–free BSA in buffer A). The supernatant was centrifuged at 12000g for 5 min at 4°C to pellet mitochondria, and the resulting pellet was washed with buffer A and finally re-suspended in ∼30 µL of buffer E solution (0.5mM EGTA in buffer A solution) to obtain 8-10 µg/µL of cardiac mitochondria. Mitochondrial ROS assay: ROS were quantified in isolated cardiac mitochondria using the Amplex Red Hydrogen Peroxide/ Peroxidase Assay Kit (Thermo Fisher) as per the instructions of the manufacturer. For each sample, 5µg of cardiac mitochondria were placed into a 96-well plate in duplicates in MiR05 containing pyruvate (5.0 mM) and malate (2.0mM) or succinate (10 mM).

Cardiomyocyte (CM) isolation: CMs were isolated with a Langendorff perfusion system with collagenase digestion.^3^ Briefly, mice were sacrificed by cervical dislocation after 10 min of heparin (1000 IU/kg, intraperitoneally, Sigma) injection. Heart and aorta were excised into the cold Ca^2+^-free buffer solution (buffer 0, 130mM NaCl, 5.4mM KCl, 0.5mM MgCl_2_, 0.33mM NaH_2_PO_4_, 0.25mM HEPES, 22mM D-Glucose, pH 7.4). The aorta was cut off below the first cranial branch and cannulated to the Langendorff apparatus without allowing air entry. Heart was perfused with Ca^2+^-free buffer solution for ∼5 min and then switched to buffer 0 containing 50 µM CaCl_2_ and type IV collagenase (200U/mL, life Technologies) for digestion with a flow rate of 2.5 mL/min. When the heart became flaccid, it was moved into buffer 0 containing 80 µM CaCl_2,_ 1.67% BSA, and 100U/mL collagenase for separating CMs. The separated cells were filtered and re-suspended in buffer 0 with 1% BSA containing either 250 µM or 500 µM or 900 µM CaCl_2_ for calcium recovery. The last cell pellet was stored in 1.8 mM CaCl_2_ in buffer 0 for cell contraction studies. CMs were studied from Sham, Sham+(L+A), OVX, OVX+(L+A) mice. Also, CMs were isolated from control mice and then treated with 0.1, 1, 10, 100, and 1000 nM oligomycin *in vitro* before measuring contraction.

Ca^2+^ transient assessment: Intracellular calcium (Ca^2+^) transients were measured using the Ca^2+^-sensitive fluo-4 AM dye (2 µM for 30 min, Life Technologies) in isolated CMs. Cells were stimulated with 1.0Hz, 10V and Ca^2+^ transients were assessed when the Ca^2+^amplitude reached steady state. Ca^2+^ release and decay rates were assessed using IonWizard 6.0 software.

RT-PCR: RNA was isolated from frozen heart tissue using TRIzol solution (Thermo Fisher Scientific Inc.) and RNeasy mini kit (Qiagen). RNA was reverse transcribed into cDNA by SuperScript II First-Strand Synthesis System (Thermo Fisher Scientific Inc.) cDNA (1.0 µg) was used for quantitative real-time PCR using Taqman probes (Applied Biosystems Inc) and primers for metabolic, heart failure and OXPHOS genes.

## References for Supplemental Methods

1. Stallcup WB, Gu Y, Dalton ND, Cedenilla M, et al. Resident fibroblast lineages mediate pressure overload-induced cardiac fibrosis. The Journal of clinical investigation. 2014;124:2921–2934. doi: 10.1172/JCI74783

2. Emde B, Heinen A, Godecke A, Bottermann K. Wheat germ agglutinin staining as a suitable method for detection and quantification of fibrosis in cardiac tissue after myocardial infarction. Eur J Histochem. 2014;58:2448. doi: 10.4081/ejh.2014.2448

3. Kohncke C, Lisewski U, Schleussner L, Gaertner C, Reichert S, Roepke TK. Isolation and Kv channel recordings in murine atrial and ventricular cardiomyocytes. Journal of visualized experiments: JoVE. 2013:e50145. doi: 10.3791/50145

## References

1. Frangogiannis NG. Cardiac fibrosis: Cell biological mechanisms, molecular pathways and therapeutic opportunities. Mol Aspects Med. 2019;65:70–99. doi: 10.1016/j.mam.2018.07.001

2. Fu M, Zhou J, Thunstrom E, Almgren T, Grote L, Bollano E, Schaufelberger M, Johansson MC, Petzold M, Swedberg K, et al. Optimizing the Management of Heart Failure With Preserved Ejection Fraction in the Elderly by Targeting Comorbidities (OPTIMIZE-HFPEF). Journal of cardiac failure. 2016;22:539–544. doi: 10.1016/j.cardfail.2016.01.011

3. Penjaskovic D, Sakac D, Dejanovic J, Zec R, Zec Petkovic N, Stojsic Milosavljevic A. Left ventricular diastolic dysfunction in patients with metabolic syndrome. Med Pregl. 2012;65:18–22.

4. Goyal P, Paul T, Almarzooq ZI, Peterson JC, Krishnan U, Swaminathan RV, Feldman DN, Wells MT, Karas MG, Sobol I, et al. Sex- and Race-Related Differences in Characteristics and Outcomes of Hospitalizations for Heart Failure With Preserved Ejection Fraction. Journal of the American Heart Association. 2017;6. doi: 10.1161/JAHA.116.003330

5. Masoudi FA, Havranek EP, Smith G, Fish RH, Steiner JF, Ordin DL, Krumholz HM. Gender, age, and heart failure with preserved left ventricular systolic function. Journal of the American College of Cardiology. 2003;41:217–223.

6. Baker L, Meldrum KK, Wang M, Sankula R, Vanam R, Raiesdana A, Tsai B, Hile K, Brown JW, Meldrum DR. The role of estrogen in cardiovascular disease. The Journal of surgical research. 2003;115:325–344.

7. Bassuk SS, Manson JE. The timing hypothesis: Do coronary risks of menopausal hormone therapy vary by age or time since menopause onset? Metabolism: clinical and experimental. 2016;65:794–803. doi: 10.1016/j.metabol.2016.01.004

8. Gupte AA, Pownall HJ, Hamilton DJ. Estrogen: an emerging regulator of insulin action and mitochondrial function. J Diabetes Res. 2015;2015:916585. doi: 10.1155/2015/916585

9. Hamilton DJ, Minze LJ, Kumar T, Cao TN, Lyon CJ, Geiger PC, Hsueh WA, Gupte AA. Estrogen receptor alpha activation enhances mitochondrial function and systemic metabolism in high-fat-fed ovariectomized mice. Physiol Rep. 2016;4. doi: 10.14814/phy2.12913

10. Morgan JP. Cellular mechanisms of diastolic dysfunction. UpToDate. 2013.

11. Phan TT, Abozguia K, Nallur Shivu G, Mahadevan G, Ahmed I, Williams L, Dwivedi G, Patel K, Steendijk P, Ashrafian H, et al. Heart failure with preserved ejection fraction is characterized by dynamic impairment of active relaxation and contraction of the left ventricle on exercise and associated with myocardial energy deficiency. Journal of the American College of Cardiology. 2009;54:402–409. doi: 10.1016/j.jacc.2009.05.012

12. Moller Petrun A, Markota A. Angiotensin II-Real-Life Use and Literature Review. Medicina (Kaunas). 2024;60. doi: 10.3390/medicina60091483

13. Murphy AM, Wong AL, Bezuhly M. Modulation of angiotensin II signaling in the prevention of fibrosis. Fibrogenesis Tissue Repair. 2015;8:7. doi: 10.1186/s13069-015-0023-z

14. Laine GA, Allen SJ. Left ventricular myocardial edema. Lymph flow, interstitial fibrosis, and cardiac function. Circ Res. 1991;68:1713–1721. doi: 10.1161/01.res.68.6.1713

15. Valentini A, Heilmann RM, Kuhne A, Biagini L, De Bellis D, Rossi G. The Renin-Angiotensin-Aldosterone System (RAAS): Beyond Cardiovascular Regulation. Vet Sci. 2025;12. doi: 10.3390/vetsci12080777

16. Schiattarella GG, Altamirano F, Tong D, French KM, Villalobos E, Kim SY, Luo X, Jiang N, May HI, Wang ZV, et al. Nitrosative stress drives heart failure with preserved ejection fraction. Nature. 2019;568:351–356. doi: 10.1038/s41586-019-1100-z

17. Hamilton DJ, Zhang A, Li S, Cao TN, Smith JA, Vedula I, Cordero-Reyes AM, Youker KA, Torre-Amione G, Gupte AA. Combination of angiotensin II and l-NG-nitroarginine methyl ester exacerbates mitochondrial dysfunction and oxidative stress to cause heart failure. American journal of physiology Heart and circulatory physiology. 2016;310:H667–680. doi: 10.1152/ajpheart.00746.2015

18. Berger JH, Shi Y, Matsuura TR, Batmanov K, Chen X, Tam K, Marshall M, Kue R, Patel J, Taing R, et al. Two-hit mouse model of heart failure with preserved ejection fraction combining diet-induced obesity and renin-mediated hypertension. Sci Rep. 2025;15:422. doi: 10.1038/s41598-024-84515-9

19. Tong D, Schiattarella GG, Jiang N, May HI, Lavandero S, Gillette TG, Hill JA. Female Sex Is Protective in a Preclinical Model of Heart Failure With Preserved Ejection Fraction. Circulation. 2019;140:1769–1771. doi: 10.1161/CIRCULATIONAHA.119.042267

20. Methawasin M, Strom J, Marino VA, Gohlke J, Muldoon J, Herrick SR, van der Pijl R, Konhilas JP, Granzier H. An ovary-intact postmenopausal HFpEF mouse model; menopause is more than just estrogen deficiency. Am J Physiol Heart Circ Physiol. 2025;328:H719–H733. doi: 10.1152/ajpheart.00575.2024

21. Mayer LP, Devine PJ, Dyer CA, Hoyer PB. The follicle-deplete mouse ovary produces androgen. Biol Reprod. 2004;71:130–138. doi: 10.1095/biolreprod.103.016113

22. Fukuta H, Goto T, Wakami K, Ohte N. Effect of renin-angiotensin system inhibitors on mortality in heart failure with preserved ejection fraction: a meta-analysis of observational cohort and randomized controlled studies. Heart Fail Rev. 2017;22:775–782. doi: 10.1007/s10741-017-9637-0

23. Strom JO, Theodorsson A, Ingberg E, Isaksson IM, Theodorsson E. Ovariectomy and 17beta-estradiol replacement in rats and mice: a visual demonstration. Journal of visualized experiments: JoVE. 2012:e4013. doi: 10.3791/4013

24. Gao S, Ho D, Vatner DE, Vatner SF. Echocardiography in Mice. Current protocols in mouse biology. 2011;1:71–83.

25. Kohncke C, Lisewski U, Schleussner L, Gaertner C, Reichert S, Roepke TK. Isolation and Kv channel recordings in murine atrial and ventricular cardiomyocytes. Journal of visualized experiments: JoVE. 2013:e50145. doi: 10.3791/50145

26. Li S, Li X, Zheng H, Xie B, Bidasee KR, Rozanski GJ. Pro-oxidant effect of transforming growth factor-beta1 mediates contractile dysfunction in rat ventricular myocytes. Cardiovasc Res. 2008;77:107–117. doi: 10.1093/cvr/cvm022

27. Benech JC, Benech N, Zambrana AI, Rauschert I, Bervejillo V, Oddone N, Damian JP. Diabetes increases stiffness of live cardiomyocytes measured by atomic force microscopy nanoindentation. American journal of physiology Cell physiology. 2014;307:C910–919. doi: 10.1152/ajpcell.00192.2013

28. Goodwin GW, Taylor CS, Taegtmeyer H. Regulation of energy metabolism of the heart during acute increase in heart work. J Biol Chem. 1998;273:29530–29539. doi: 10.1074/jbc.273.45.29530

29. Rijzewijk LJ, van der Meer RW, Lamb HJ, de Jong HW, Lubberink M, Romijn JA, Bax JJ, de Roos A, Twisk JW, Heine RJ, et al. Altered myocardial substrate metabolism and decreased diastolic function in nonischemic human diabetic cardiomyopathy: studies with cardiac positron emission tomography and magnetic resonance imaging. Journal of the American College of Cardiology. 2009;54:1524–1532. doi: 10.1016/j.jacc.2009.04.074

30. Thorn SL, deKemp RA, Dumouchel T, Klein R, Renaud JM, Wells RG, Gollob MH, Beanlands RS, DaSilva JN. Repeatable noninvasive measurement of mouse myocardial glucose uptake with 18F-FDG: evaluation of tracer kinetics in a type 1 diabetes model. J Nucl Med. 2013;54:1637–1644. doi: 10.2967/jnumed.112.110114

31. Baessler A, Strack C, Rousseva E, Wagner F, Bruxmeier J, Schmiedel M, Riegger G, Lahmann C, Loew T, Schmitz G, et al. Growth-differentiation factor-15 improves reclassification for the diagnosis of heart failure with normal ejection fraction in morbid obesity. Eur J Heart Fail. 2012;14:1240–1248. doi: 10.1093/eurjhf/hfs116

32. Duerrschmid C, Trial J, Wang Y, Entman ML, Haudek SB. Tumor necrosis factor: a mechanistic link between angiotensin-II-induced cardiac inflammation and fibrosis. Circ Heart Fail. 2015;8:352–361. doi: 10.1161/CIRCHEARTFAILURE.114.001893

33. Daou D, Tong D, Schiattarella GG, Gillette TG, Hill JA. What Is Cardiometabolic HFpEF and How Can We Study it Preclinically? JACC Basic Transl Sci. 2025;10:101295. doi: 10.1016/j.jacbts.2025.04.009

34. Withaar C, Meems LMG, Markousis-Mavrogenis G, Boogerd CJ, Silljé HHW, Schouten EM, Dokter MM, Voors AA, Westenbrink BD, Lam CSP, et al. The effects of liraglutide and dapagliflozin on cardiac function and structure in a multi-hit mouse model of heart failure with preserved ejection fraction. Cardiovasc Res. 2021;117:2108–2124. doi: 10.1093/cvr/cvaa256

35. Sharma K, Kass DA. Heart failure with preserved ejection fraction: mechanisms, clinical features, and therapies. Circulation research. 2014;115:79–96. doi: 10.1161/CIRCRESAHA.115.302922

36. Bazgir F, Nau J, Nakhaei-Rad S, Amin E, Wolf MJ, Saucerman JJ, Lorenz K, Ahmadian MR. The Microenvironment of the Pathogenesis of Cardiac Hypertrophy. Cells. 2023;12. doi: 10.3390/cells12131780

37. Hamdani N, Franssen C, Lourenco A, Falcao-Pires I, Fontoura D, Leite S, Plettig L, Lopez B, Ottenheijm CA, Becher PM, et al. Myocardial titin hypophosphorylation importantly contributes to heart failure with preserved ejection fraction in a rat metabolic risk model. Circ Heart Fail. 2013;6:1239–1249. doi: 10.1161/CIRCHEARTFAILURE.113.000539

38. Peana D, Domeier TL. Cardiomyocyte Ca(2+) homeostasis as a therapeutic target in heart failure with reduced and preserved ejection fraction. Current opinion in pharmacology. 2017;33:17–26. doi: 10.1016/j.coph.2017.03.005

39. Ingwall JS, Weiss RG. Is the failing heart energy starved? On using chemical energy to support cardiac function. Circulation research. 2004;95:135–145. doi: 10.1161/01.RES.0000137170.41939.d9

40. Senni M, Paulus WJ, Gavazzi A, Fraser AG, Diez J, Solomon SD, Smiseth OA, Guazzi M, Lam CS, Maggioni AP, et al. New strategies for heart failure with preserved ejection fraction: the importance of targeted therapies for heart failure phenotypes. European heart journal. 2014;35:2797–2815. doi: 10.1093/eurheartj/ehu204

41. Koves TR, Ussher JR, Noland RC, Slentz D, Mosedale M, Ilkayeva O, Bain J, Stevens R, Dyck JR, Newgard CB, et al. Mitochondrial overload and incomplete fatty acid oxidation contribute to skeletal muscle insulin resistance. Cell Metab. 2008;7:45–56.

42. Pabel S, Wagner S, Bollenberg H, Bengel P, Kovacs A, Schach C, Tirilomis P, Mustroph J, Renner A, Gummert J, et al. Empagliflozin directly improves diastolic function in human heart failure. Eur J Heart Fail. 2018;20:1690–1700. doi: 10.1002/ejhf.1328

